# Unravelling the shared genetic mechanisms underlying 18 autoimmune diseases using a systems approach

**DOI:** 10.1101/2021.03.28.437437

**Authors:** Sreemol Gokuladhas, William Schierding, Evgeniia Golovina, Tayaza Fadason, Justin O’Sullivan

## Abstract

Autoimmune diseases (AiDs) are complex heterogeneous diseases characterized by hyperactive immune responses against self. Genome-wide association studies have identified thousands of single nucleotide polymorphisms (SNPs) associated with several AiDs. While these studies have identified a handful of pleiotropic loci that confer risk to multiple AiDs, they lack the power to detect shared genetic factors residing outside of these loci. Here, we integrated chromatin contact, expression quantitative trait loci and protein-protein interaction (PPI) data to identify genes that are regulated by both pleiotropic and non-pleiotropic SNPs. The PPI analysis revealed complex interactions between the shared and disease-specific genes. Furthermore, pathway enrichment analysis demonstrated that the shared genes co-occur with disease-specific genes within the same biological pathways. In conclusion, our results are consistent with the hypothesis that genetic risk loci associated with multiple AiDs converge on a core set of biological processes that potentially contribute to the emergence of polyautoimmunity.

## Introduction

Autoimmune diseases (AiDs) are chronic conditions that arise when there is an abnormal immune response that targets functioning organs. Many AiDs share clinical symptoms and immunopathological mechanisms (Anaya, 2012). For instance, it has been shown that patients with the most common AiDs such as multiple sclerosis (MS), type I diabetes (TID), rheumatoid arthritis (RA), and systemic lupus erythematosus (SLE) are at higher risk of polyautoimmunity (Bao et al., 2019; Ordoñez-Cañizares et al., 2020; Ramagopalan, Dyment, & Ebers, 2008). It is likely that environmental factors impact on the shared immunopathological mechanisms to trigger polyautoimmunity. On the other hand, there is evidence for a genetic contribution to AiD development that is supported by higher concordance rates in monozygotic twins, a relative increase in the risk of disease in dizygotic twins (Bogdanos et al., 2012), and the coexistence of AiDs within families and/or individuals (Mäkimattila, Harjutsalo, Forsblom, & Groop, 2020; Simon et al., 2020, 2017; Somers, Thomas, Smeeth, & Hall, 2006). We hypothesize that the effects of AiD associated genetic variants converge on biological pathways that increase risk through downstream functional impacts.

The major histocompatibility complex (MHC) locus provides the greatest genetic risk factor for AiD development and is an obvious common link between AiDs (Matzaraki, Kumar, Wijmenga, & Zhernakova, 2017). In addition to the MHC locus, non-HLA genes such as *CTLA4*, *PTPN22*, and *TNF* have also been associated with multiple AiDs (Serrano, Millan, & Páez, 2006). Furthermore, genome-wide association studies (GWAS) have identified thousands of single nucleotide polymorphisms (SNPs) across the human genome that are associated with an increased risk of developing AiD. The AiDs-associated GWAS SNPs are typically inter-genic and unique to one, or small set of AiDs (Lettre & Rioux, 2008). Given the phenotypic similarities between the AiDs, it is however possible that combined analyses may reveal patterns of shared genetic and pathological etiology. Consistent with this, a cross-disease Immunochip SNP meta-analysis identified novel pleiotropic risk loci that represent complex comorbidity from patients with seronegative immune phenotypes (Ellinghaus et al., 2016).

Trait-associated SNPs have been shown to be more likely to mark loci that are expression quantitative trait loci (eQTL)(Nicolae et al., 2010). In this study, we have concurrently investigated SNPs that were independently associated with 18 AiDs to identify their transcriptional regulatory activity (*i.e*., as eQTLs), using an *in silico* method (CoDeS3D) that combines different levels of empirical evidence (Fadason, Schierding, Lumley, & O’Sullivan, 2018). We further identified the target genes of the eQTLs and analysed the functional and physical interactions among the proteins they encode. Using a modularity-based community detection method, we extracted the functional modules from the protein-protein interactions. Functional enrichment analysis of the modules provided a measure of how genetically related AiD-associated genes contribute to increasing the risk of developing polyautoimmune conditions.

## Methods

### Identification of the target genes of autoimmune disease-associated SNPs

SNPs associated (p≤5×10^−6^) with 18 autoimmune diseases [alopecia areata (ALO), ankylosing spondylitis (AS), celiac disease (CED), Crohn’s disease (CRD), eosinophilic esophagitis (EE), Graves’ disease (GRD), juvenile idiopathic arthritis (JIA), multiple sclerosis (MS), primary biliary cirrhosis (PBC), psoriatic arthritis (PA), psoriasis (PSO), rheumatoid arthritis (RA), Sjogren’s syndrome (SJS), systemic lupus erythematosus (SLE), systemic scleroderma/sclerosis (SSC), type-I diabetes (T1D), ulcerative colitis (ULC), and vitiligo (VIT)] were retrieved from the GWAS catalog (https://www.ebi.ac.uk/gwas; on 30 April 2020) (Supplementary data 1). The SNPs associated with each disease were analysed separately through a python-based bioinformatics algorithm (CoDeS3D) (Fadason et al., 2018) to identify which SNPs acted as expression Quantitative Trait Loci (eQTLs) and to identify their target genes. Firstly, CoDeS3D uses Hi-C chromatin contact data derived from 70 cell lines and primary tissues (Supplementary data 2) to identify target genes that are spatially interacting with the SNPs. Secondly, eQTL data from 49 human tissues (GTEx V8) (Aguet et al., 2020)) were used to identify the SNPs (eQTLs) that are associated with the expression changes of their target genes (eGenes). Lastly, false positive associations were controlled using a multiple testing correction (Benjamini-Hochberg False Discovery Rate (FDR < 0.05)). Chromosome positions of SNPs and genes are reported according to the Human reference genome GRCh38/hg38 assembly.

### Construction of the autoimmune disease network using protein-protein interaction (PPI) data

The python ‘networkx’ library was used to construct the autoimmune disease network in two steps: (i) A reference PPI network (ref-PPIN) was constructed using data downloaded from STRING v11.0 (Szklarczyk et al., 2019). Only protein pairs with no self-links and a high-confidence score (combined score > 0.7) were retained, yielding a reference network with 16758 proteins (nodes) and 411585 interactions (edges). (ii) All genes whose expression changes were correlated with the SNPs from one or more of the 18 autoimmune diseases were analyzed to determine if they were involved in PPIs within the ref-PPIN. The resulting autoimmune PPI network (Ai-PPIN) consisted of 2925 proteins and 19173 interactions. Cytoscape (version 3.8.2) was used for PPI network visualization.

### Identification of modules from the autoimmune PPI network (Ai-PPIN)

Functional modules can be defined as either: a) a stable protein complex; or b) a set of transiently interacting proteins that together act to accomplish a specific biological function. Here, we extracted the functional modules from the Ai-PPIN using the Louvain module detection algorithm (Blondel, Guillaume, Lambiotte, & Lefebvre, 2008). The Louvain algorithm identifies functional modules by optimizing the modularity (Q) of the network. For an undirected graph *G=(V, E)* with *V* number of nodes and *E* number of edges, Q is defined as (Dugué & Perez, 2015),

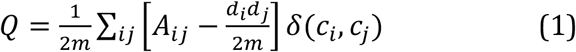

where *m* is the number of edges (*E*) of *G*, *A_ij_* represents the weight of the edge between nodes *i* and *j*, *d_i_* and *d_j_* are degrees of node *i* and *j*, *c_i_* and *c_j_* are the communities to which *i* and *j* belong, and δ- function for which *δ*(*c_i_*, *c_j_*) equals 1 if *c_i_*=*c_j_*, and 0 if *c_i_*≠ *c_j_*. The communities or the functional modules are found by maximizing the Q function in an iterative manner. In the initial stage, all nodes in the network are considered as independent modules and the algorithm progressively combines two modules that increase the Q of the resulting network. Combining nodes and modules continues until there is no further increase in the Q of the network. The Louvain module detection algorithm has previously been proposed to be the best method to find modules within the human PPI network (Rahiminejad, Maurya, & Subramaniam, 2019).

The qs-test was used to evaluate the significance of modules according to the quality function (*q*) and size (*s*) of the module. A module, M, is deemed significant if its quality function, *q_M_*(modularity), is larger than those for detected modules of the same size *s_M_* in randomized networks (Kojaku & Masuda, 2018). The size function is calculated by summing the degrees of nodes in a module.

### Identification of central genes within the functional modules

In network theory, the centrality of a node measures its relative importance within the network. We regarded each module identified from Ai-PPIN as an individual network and identified central nodes using three centrality measures: degree, closeness, and eigenvector. The python package “networkx” was used for centrality analysis.

*Degree centrality (DC)*. The DC indicates the number of direct neighbors of a node. The DC of a node *i* is defined as,

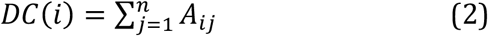

where *A* is the adjacency matrix, and *n* is the total number of nodes in a graph (*G*). DC values are normalized by dividing them by the maximum possible degree (*n − 1*), where *n* is the number of nodes in *G*.

*Closeness centrality (CC)*. The CC is the reciprocal of average shortest path distance between a node *i* and all other reachable nodes in the network. CC of a node *i* is defined as,

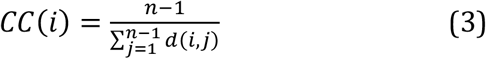

where *d*(*i, j*) is the shortest path distance between *i* and *j*, and *n* is the number of nodes that can reach *i*.

*Eigenvector centrality (EC)*. The EC computes the centrality of a node based on the centrality of its neighbours. EC measures the influence of a node on the connectivity of the network. EC of a node *i* is defined as,

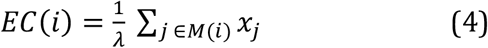

where *M*(*i*) is a set of neighbours of *i*, *λ* is the largest eigenvalue of *A*(adjacency matrix). If a node is connected to other well-connected nodes in the PPI, it will have the maximum EC value.

We sorted the proteins in decreasing order according to their degree, closeness and eigenvector centrality scores and selected the top 10% of proteins from each group. We defined the proteins that are present in common across all three groups as central.

### Functional annotation of the modules

Pathway and GO enrichment analyses were performed (R package g:profiler (version 2_0.1.9) (Raudvere et al., 2019)) on every module detected from Ai-PPIN to identify significantly enriched pathways and biological processes terms (false discovery rate correction threshold of 0.05). Kyoto Encyclopedia of Genes and Genomes (KEGG) pathways (accessed 10-October-2020) and gene ontology (GO) biological processes (accessed 20-January-2021) terms were used as the reference libraries in these analyses. DGIdb version 3.0 (Cotto et al., 2018) was used to identify potential drug interactions with the eGenes.

## Results

### An overview of the gene regulatory network of the AiDs

The SNP-gene regulatory network encompassing 2065 eQTLs (70% of the total input SNPs (N=2953)) and 4789 eGenes across 18 diseases (Supplementary data 3) was identified using CoDeS3D (Fadason et al., 2018) (Figure 1A). The eQTLs and eGenes are hereafter referred to as “SNPs” and “genes” for simplicity. Almost all SNPs (N=1879; 91%) are non-pleiotropic (*i*.e., associated with only one AiD). There are pleiotropic SNPs (N=186; 9%) implicated in two or more AiDs (Figure 1B), where two or more GWAS on different diseases independently identified the same SNP. Of these, approximately one-third of the pleiotropic SNPs (N=60; 32.3%) were associated only between CRD and ULC. The remaining 126 (67.7%) were shared between two to five disease conditions (Supplementary data 4). Together, the pleiotropic SNPs are associated with the expression levels of 833 (17.4%) genes. A small proportion of genes (N=225; 4.7%) are regulated only by pleiotropic SNPs (figure 1B, (i) termed as “identical genes”), 608 genes (12.7%) regulated by both pleiotropic and non-pleiotropic SNPs and 889 genes (18.6%) regulated by >2 non-pleiotropic SNPs associated with different AiDs (figure 1B, (ii) termed as “shared genes”). However, the vast majority of the genes (N=3067; 64%) were unique to each disease condition (figure 1B, (iii) termed as “disease-specific”). These observations are consistent with the existence of a shared genetic architecture between autoimmune diseases that is primarily manifested by the disease-specific genetic mechanisms.

The 2065 SNPs identified from the 18 AiDs were connected to the 4789 genes via 9183 cis and 5414 trans regulatory interactions across 49 tissues (Supplementary data 3). However, only 40% (N=1914) of the genes were regulated by cis-SNPs and 52% (N=2498) were regulated by trans-SNPs (Figure 1C). The vast majority of trans-genes 84% (N=2100) were identified in only one of the 49 tissues analyzed. (Figure 1D). This observation suggests that the impacts of the AiD associated SNPs are largely tissue-specific in nature.

**Figure 1.**
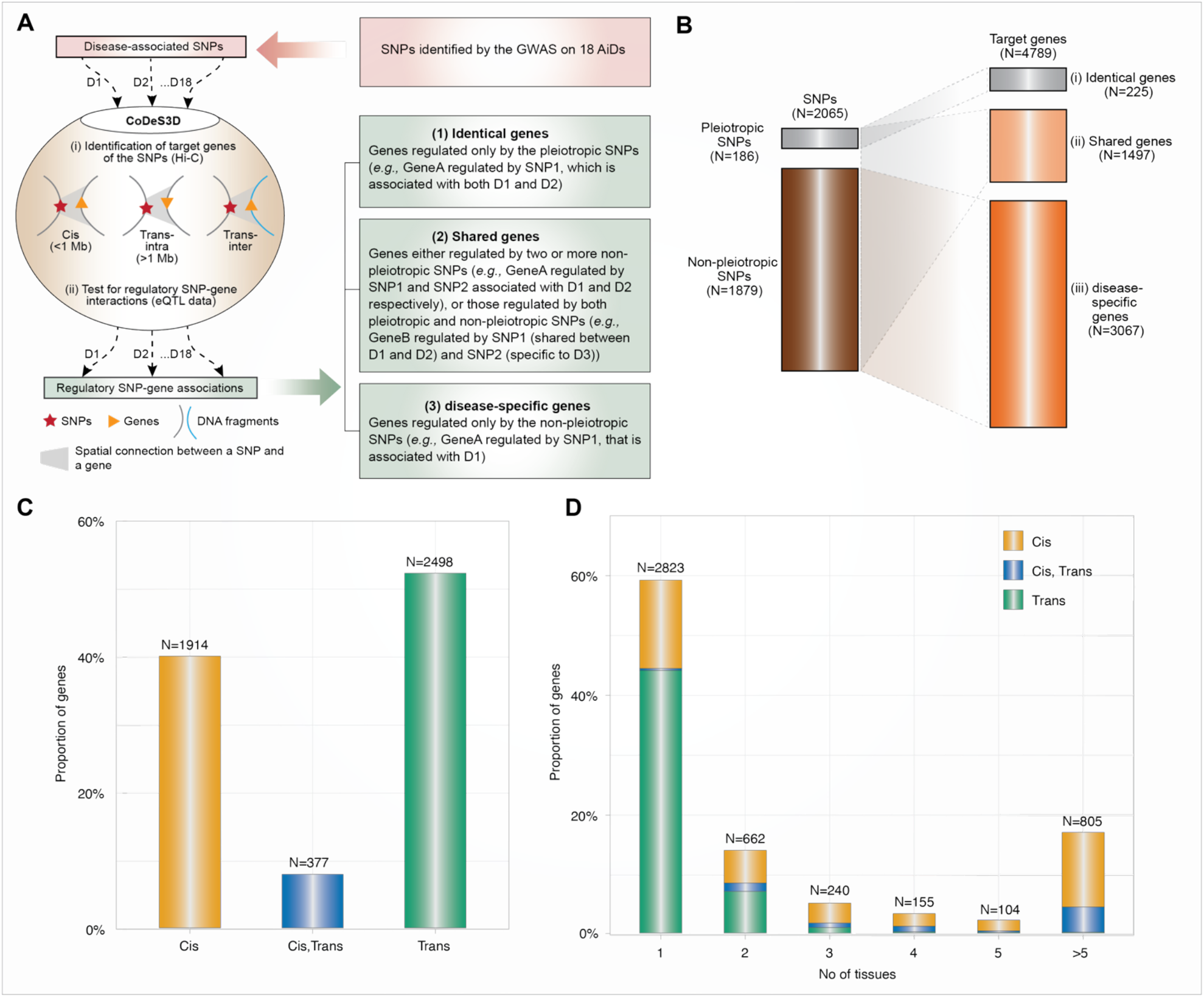
Global overview of the genetic architecture of AIDs. **(A)**SNPs associated with each of 18 AiDs (D1 to D18) were analyzed through the CoDeS3D algorithm (Fadason et al., 2018). Briefly: (i) genes that are in physical contact with the SNPs (cis - located within 1 Mb distance, trans-intrachromosomal- located on the same chromosome but more than 1 Mb apart, and trans-interchromosomal - located on the different chromosomes) within the three-dimensional organization of the nucleus are identified; and (ii) SNP-gene pairs are queried through GTEx to identify those that overlap eQTL-eGene correlations. Lastly, the regulatory SNP-gene associations identified for each of 18 AiDs were consolidated to identify the genes (1), associated with pleiotropic SNPs only, (2) associated with pleiotropic & non-pleiotropic SNPs, or >2 non-pleiotropic SNPs associated with different AiDs and, (3) associated with non-pleiotropic SNPs only. **(B)** Summary of pleiotropic and non-pleiotropic SNPs (left) and their target genes (right) across 18 AiDs by proportion. Dotted lines indicate associations between categories of SNPs and genes. **(C)** The proportion of genes regulated in cis, trans (inter- and/or intra-chromosomal), or both cis and trans by the SNPs across 18 AiDs. **(D)** Trans-regulatory connections were enriched in single tissue. Proportion of genes was calculated as percentage total genes.

### AiD associated genes organize into highly modular communities

We constructed an autoimmune protein-protein interaction network (Ai-PPIN) for the proteins encoded by the genes we identified. The schematic representation of the network analysis is presented (Figure 2A). Non-coding genes and those with missing entrez gene identifiers were filtered from the PPI analysis, resulting in a set of 4253 genes, of which Ai-PPIN contained the protein products of 2925 genes (Supplementary data 5 Table 1) and 19173 interactions (Supplementary data 5 Table 2).

**Figure 2.**
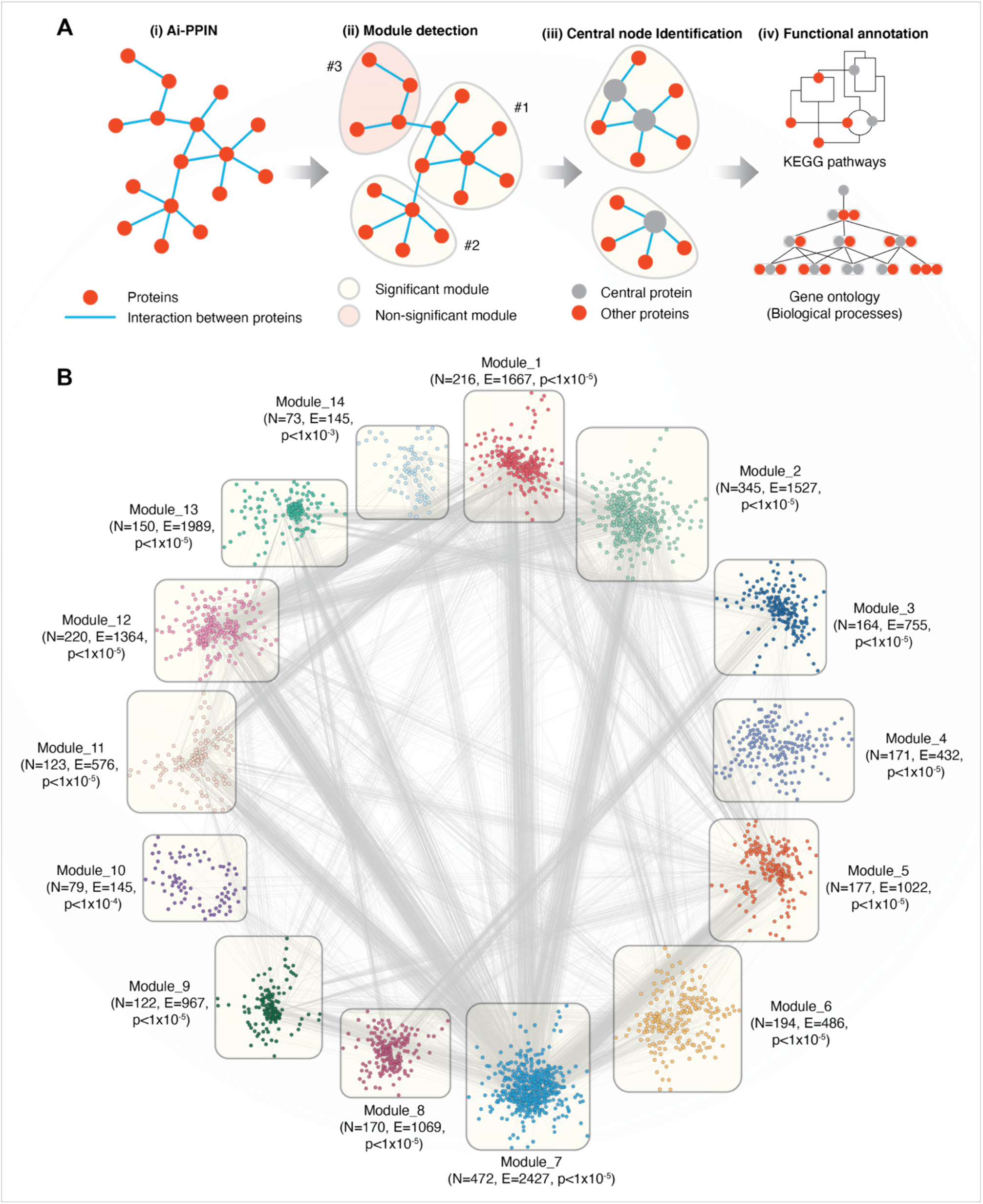
Overview of the functional modules identified from Ai-PPIN. **(A).** Schematic representation of the Ai-PPIN module analysis. The Louvain community detection algorithm (Blondel et al., 2008) was applied to detect communities/modules within the Ai-PPIN network. Statistically significant (qs-test) (Kojaku & Masuda, 2018) modules (yellow bubble) were identified by comparison with modules from 10000 random networks. Non-significant modules (red bubble) were excluded from further analysis. Functional enrichment analyses using KEGG pathways and GO:BP (gene ontology biological process terms) were performed to identify the biological functions enriched within each module. **(B)** The Ai-PPIN contains fourteen significant modules. In each module, the nodes represent proteins. The lines connecting the nodes represent interactions between proteins. N and E denotes the number of nodes and edges present in each module respectively. The p-value denotes the statistical significance of the modules (qs-test) (Kojaku & Masuda, 2018). Cytoscape (version 3.8.2) was used for visualization of the network.

It is established that within a biological network, disease-associated genes are likely to form modules that are important for the cellular processes underlying disease pathogenesis (Sharma et al., 2014). We identified network modules using the Louvain community detection algorithm (Blondel et al., 2008) and tested their statistical significance against 10000 randomly generated networks using the qs-test (Kojaku & Masuda, 2018). The Louvain algorithm detected 81 potential modules from the network, of which 14 were statistically significant. These 14 significant modules contained between 73 to 472 proteins each and accounted for 2676 of the proteins in the Ai-PPIN (Figure 2B, Supplementary data 6). The remaining 249 proteins assembled into 67 non-significant modules were excluded from the analysis. As expected, the gene products encoded by the HLA genes exhibited high interaction and were organized into a single module (Module 1). The aggregation of proteins into distinct communities within the Ai-PPIN suggests a high tendency of AiD associated proteins to physically or functionally interact to perform the intended cellular function.

We annotated the functions of the modules using KEGG pathways enrichment analysis. According to the top 5 significantly enriched pathways, each module is classified with distinct biological functions. For instance, Module 1 is enriched for proteins involved in pathways related to immune system and immune diseases; Module 11 is enriched for endocytosis and infectious disease related pathways; Module 3, 8 and 13 for genetic information processing pathways (e.g., RNA degradation, spliceosome, Ubiquitin mediated proteolysis), Module 4, 10 and 14 for distinct metabolic pathways (Supplementary data 7). Each functional module exhibits functional heterogeneity, meaning that they are involved in diverse biological functions. Functional heterogeneity of the modules suggest that they may consist of one or more transiently interacting protein complexes (Li, Wu, Wang, & Pan, 2012), which also reveal a potential link between apparently unrelated biological processes.

### Shared genes display predominant role in AiD modules

Altogether, the significant modules identified within the Ai-PPIN network are composed of approximately 30% shared, 65% disease-specific, and 4% identical proteins. Module 14 is an exception as it does not contain any protein encoded by identical genes. Within each module, at least 12 AiDs were represented by disease-specific proteins. Notably, all 18 AIDs were represented by disease-specific proteins in Modules 2, 3, and 12. This is consistent with the hypothesis that interactions between multiple AiD associated proteins may contribute to co-morbid features. Remarkably, the proportion of shared proteins is considerably larger than those of the disease-specific or identical proteins in all 14 modules (Figure 3A). KEGG pathway analysis identified that 34% (18.5714±7.764) of proteins that are enriched within the top 5 biological pathways are shared between multiple AiDs (Figure 3B). Moreover, the shared proteins are also essential to the modules as confirmed by the centrality analysis (Supplementary data 8). Notably, more than 50% of the proteins representing central nodes in Module 1 (enriched for immune pathways) and Module 4 (enriched for metabolic pathways) are shared between AiDs (Figure 3C). The co-occurrence of shared proteins in central positions within the pathways containing disease-specific proteins might contribute to the risk of developing comorbid conditions.

**Figure 3.**
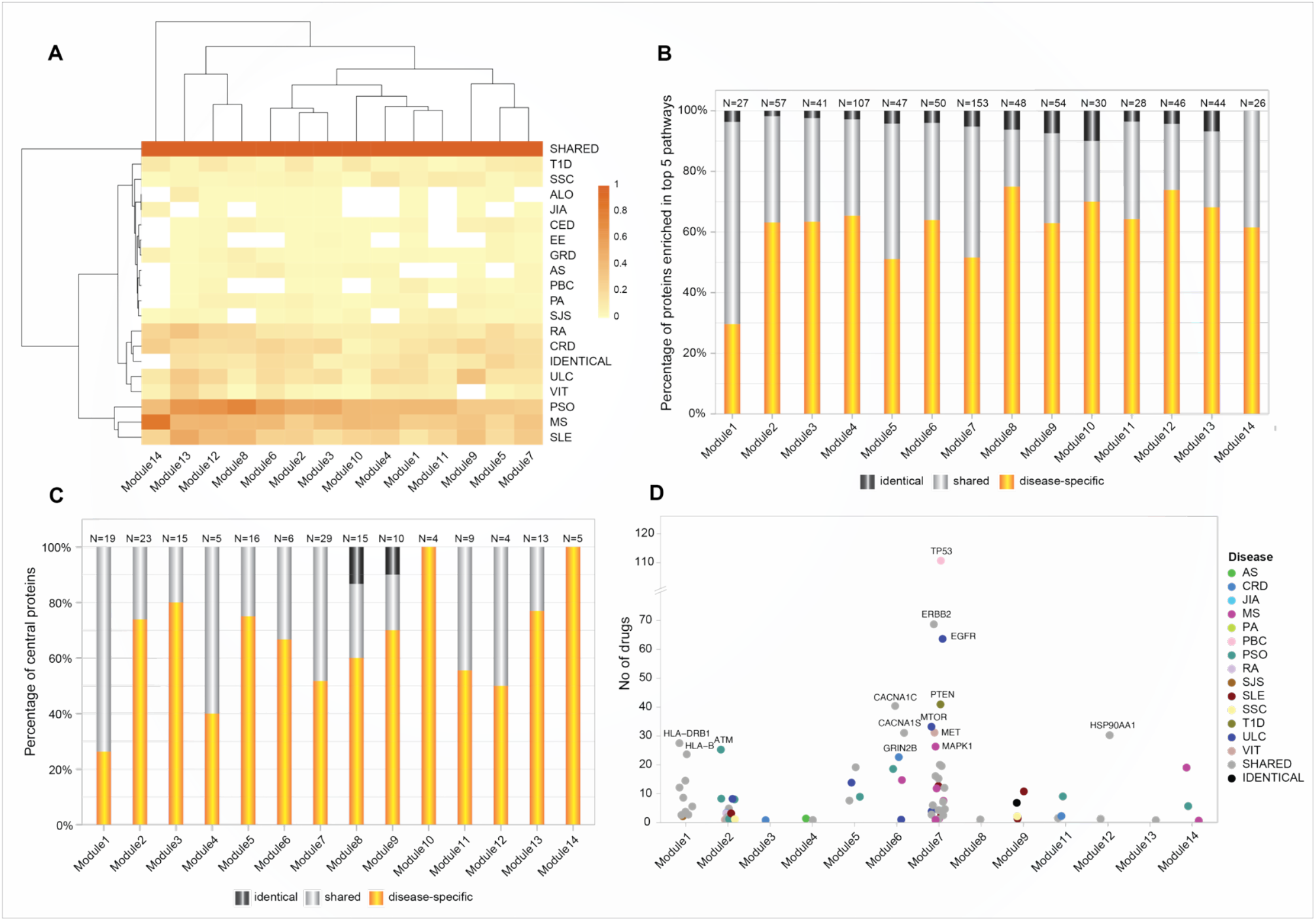
Shared genes display predominant role in AiD modules. **(A)** Heatmap of proportion of genes/proteins from each AiD that were attributed to modules 1-14. Dark shaded square indicates higher proportions of proteins. **(B)** The proportion of shared, disease-specific and identical proteins present in the top 5 enriched biological pathways (KEGG), by module. **(C)** The proportion of disease-specific, shared and identical proteins that constitutes central nodes within each module. **(D)** The central proteins in 13 modules are targeted by FDA approved drugs, of which 45% proteins are shared between diseases. Proteins that are targeted by more than 20 drugs are labeled.

DGIdb analysis determined that 80 of 173 (about 46%; Supplementary data 9 Table 1) of the central proteins across the 14 modules have known drug targets with 45% of the druggable proteins being shared between AiDs (Figure 3D; Supplementary data 9 Table 2). These proportions are much greater than the proportion of GENCODE genes with known drug targets (4807 out of 54592, 9%), which informs the pharmacological value of the central and shared proteins, respectively.

### Human leukocyte antigen (HLA) genes are central to immune function rich module

Genetic risk for autoimmune diseases including T1D, CED, autoimmune thyroid disease, SJS, SLE, RA, MS, and autoimmune hepatitis (Cruz-Tapias et al., 2012; Fridkis-Hareli, 2008) has been previously attributed to variants within the MHC region. Consistent with this, we observed that proteins encoded by the MHC region genes interact with other non-MHC gene products to form the densely connected Module 1 (Figure 4A) (clustering coefficient=0.586; indicates greater connectivity of the neighborhood of the nodes). Module 1 contains disease-specific proteins (60%), associated with 17 AiDs, shared (34%) and identical proteins (6%; Supplementary data 10 Table 1). Gene ontology analysis revealed that the 199 proteins located within Module 1 are overrepresented in 677 biological processes (Supplementary data 10 Table 2), including significantly enriched terms related to cellular transport, localization and the immune system associated functions (Figure 4B). KEGG pathway enrichment analysis confirmed significant enrichment in pathways that are predominantly linked to immune system, immune diseases, and infectious diseases (Figure 4C; Supplementary data 10 Table 3). Centrality analysis identified that the HLA class I and II proteins and six other proteins (CAPZB, CAPZA1, CAPZA2, DCTN2, ACTR1A, and DYNC1I1) as being most essential within Module 1 (Figure 4A). Notably, the significantly enriched biological process terms (N=29 of top 30) and pathways (N=33 of 44) contained shared proteins that were central to the module (Figure 4B and 4C; Supplementary data 10 Table 4 and 5). Similarly, the expression of transcripts from the HLA-DQA2, HLA-DRB1, HLA-DQB1, HLA-DRA, HLA-DRB5, HLA-G, and HLA-C genes is altered by SNPs associated with between 11 to 16 AiDs (Figure 4A, Supplementary data 10 Table 6). These observations are consistent with the central role(s) for HLA encoded genes in the pathogenesis of AIDs. The interactions involving HLA genes, that are highly influenced by the epistatic interaction of multiple disease-specific SNPs, may potentially modulate the biological processes or pathways related to immune system response and functions.

**Figure 4.**
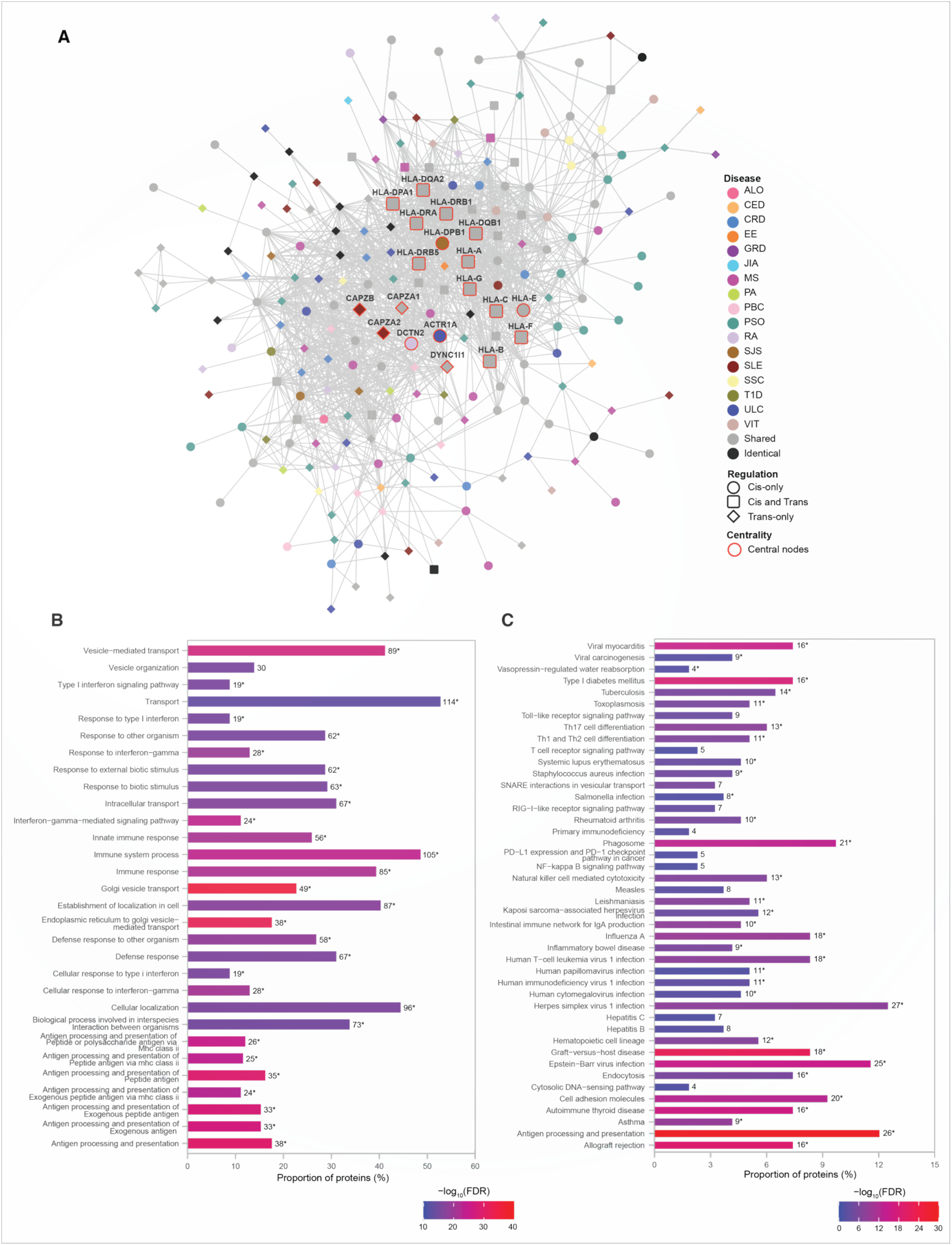
HLA genes are central to immune function rich module. **(A)** Network representation of Module 1. The color of the nodes denotes the disease with which the protein is associated. Node shape indicates if the SNP acts locally (cis - circle), distally (trans - diamond), or both (cis and trans – rounded square) on the genes encoding proteins. Central nodes are highlighted in red borders and labelled. Cytoscape (version 3.8.2) was used for visualization of the module. **(B)** Relatively greater proportions of proteins (>40%) in the Module 1 are enriched for transport, localization and immune processes. The top 30 enrichment results are shown (FDR≤6.01e-14) **(C)** KEGG pathway enrichment analysis identified enrichment in immune related pathways (FDR<0.05). In **(B)** and **(C)**, the numbers on top of each bar denote the number of proteins enriched for that term/pathway, and the asterisk denotes that the term/pathway is also enriched for shared proteins that are central to the network (Supplementary data 10 Table 4 and 5).

### Non-HLA proteins organize into a module enriched for immune responses

Module 5 consists of 177 proteins (Supplementary data 11 Table 1), 59% of which are associated with one of 16 AiDs, with a clustering coefficient of 0.568. In contrast to Module 1, three-fourth (12 out of 16; 75%) of the central proteins within module 5 is disease-specific (Figure 5A). The central proteins that are shared between conditions are associated with two to six AiDs. For example, PLAU is shared between CRD (rs2227551, rs2227564), MS (rs2688608), and PSO (rs2675662); ITGAM is shared between GRD (rs57348955), PSO (rs12445568, rs10782001, rs13708) and SLE (rs11150610); RAP1A is shared between CRD (rs488200) and PSO (rs11121129); and *ATP8B4* is targeted by the pleiotropic SNPs rs12946510, rs12946510, rs12946510 - associated with CRD, MS, and ULC; rs2305480, rs2305480 -associated with RA and ULC; and non-pleiotropic SNPs- rs883770 (SSC), and rs2290400 (TID). The proteins within Module 5 are significantly enriched for ontological terms including immune response and transport (Supplementary data 11 Table 2) and biological pathways related to cellular signaling, infectious diseases and immune system (Supplementary data 11 Table 3). Furthermore, the shared central proteins are involved in the biological processes (N=29 of top 30) predominantly linked to immune responses (Figure 5B; Supplementary data 11 Table 4) and KEGG pathways (N=10 of 19) including those linked to immune processes such as complement and coagulation cascades, hematopoietic cell lineage and leukocyte transendothelial migration (Figure 5C; Supplementary data 11 Table 5). The enrichment of proteins in Module 5 for the immune system related processes can lead to speculation that non-HLA loci may contribute to the AiD pathology by modulating alternate immune response pathways.

**Figure 5.**
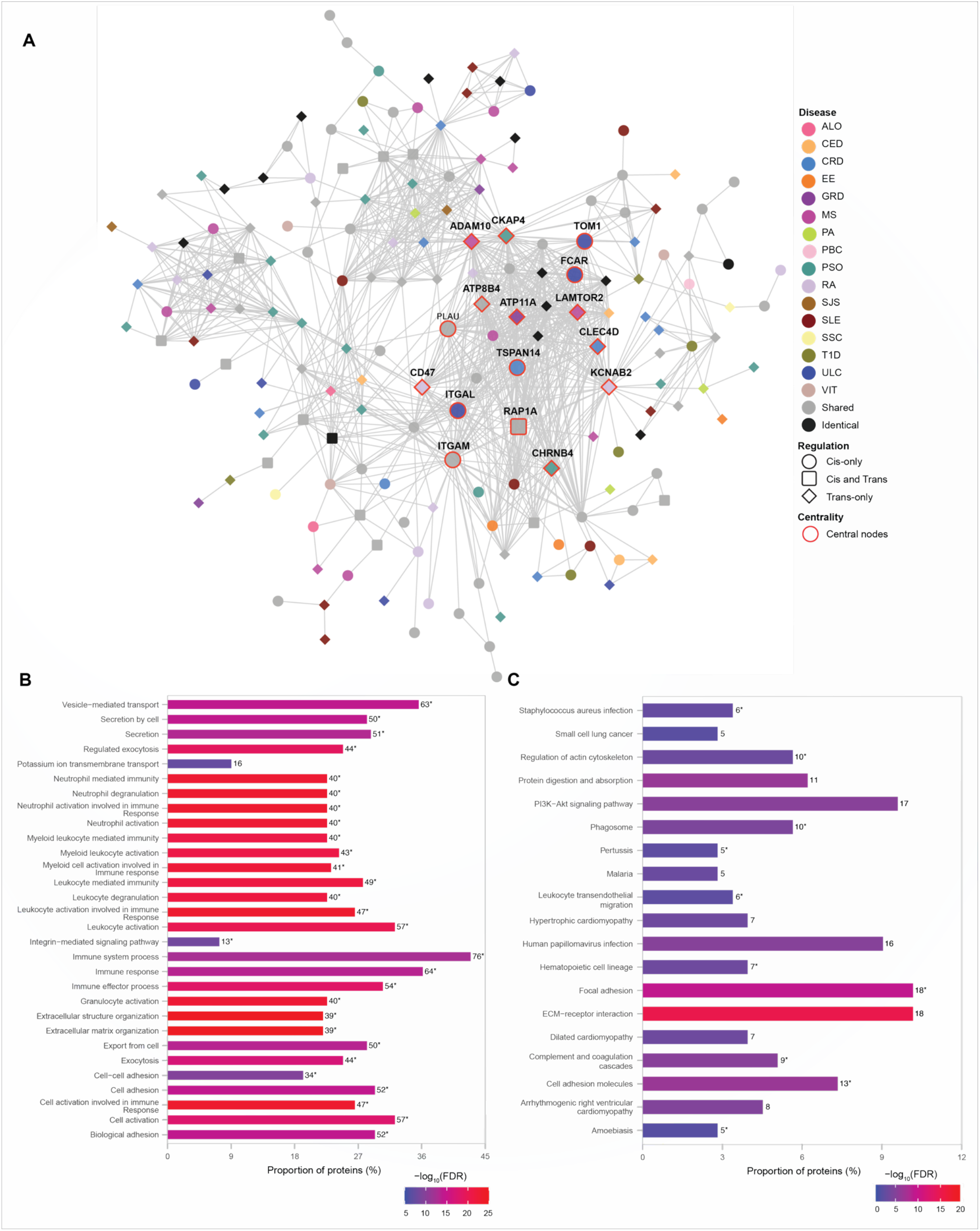
Non-HLA proteins organize into a module enriched for immune responses. **(A)** Network representation of Module 5. The color of the nodes denotes the disease with which the protein is associated. Node shape indicates if the SNP acts locally (cis - circle), distally (trans - diamond), or both (cis and trans – rounded square) on the genes encoding proteins. Central nodes are highlighted in red borders and labelled. Cytoscape (version 3.8.2) was used for visualization of the module. **(B)** Module 5 is highly enriched for immune processes. The top 30 enrichment results are shown (FDR≤5.6e-09) **(C)** KEGG pathway enrichment results with FDR<0.05 is shown. In **(B)** and **(C)**, the numbers, to the right of each bar, denote the number of proteins enriched for that term or pathway. The asterisk designates terms or pathways that were also enriched for shared proteins that are central to the network (Supplementary data 11 Tables 4 and 5 respectively).

### The largest network module is enriched for cellular signalling and cancer pathways

Module 7 is the largest (N=472 proteins) functional module, with the clustering coefficient of 0.425, identified from the Ai-PPIN network. As observed for modules 1 and 5, the bulk of the proteins within module 7 is encoded by disease-specific genes (281: 163: 28, disease-specific: shared: identical; Supplementary data 12 Table 1). As observed for Module 1, a large proportion (48%; N=14 of 29) of the central nodes within Module 7 is shared proteins. However, some disease-specific proteins are also central to this cluster. For example, the transcript levels of tumor-suppressor gene *TP53* are associated only with a PBC-associated SNP (rs12708715). However, TP53 interacts with 62 other proteins (42 and 20 encoded by disease-specific and shared, respectively) within Module 7. Transcript levels of an additional twelve cancer-related genes (*i.e., HRAS, ERBB2, STAT3, RHOA, SYK, MAP2K1, LYN, PRKCB, NFKB1, MAPK3, IL2RA*, and *GRB2*; human protein atlas) are associated with SNPs from more than two AiDs and also highly interconnected with other genes in Module 7. GO analysis identified enrichment for biological process terms associated with system-wide regulatory activities (Figure 6b; Supplementary data 12 Table 2). Similarly, KEGG pathway analyses indicated that Module 7 is enriched for proteins that are involved in axon guidance, immune function, cellular signaling, cancer, apoptosis, and infectious diseases (Figure 6c; Supplementary data 12 Table 3). Collectively, these results indicate that the impacts of proteins within Module 7 is not only limited to specific cellular mechanisms but may disrupt wider processes during the course of development of a disease. Moreover, Module 7 provides a potential mechanism for observed increases in multimorbidity between AiDs and certain forms of cancer (Hai-long Wang, Zhou, Zhu, Yang, & Hua, 2018).

**Figure 6.**
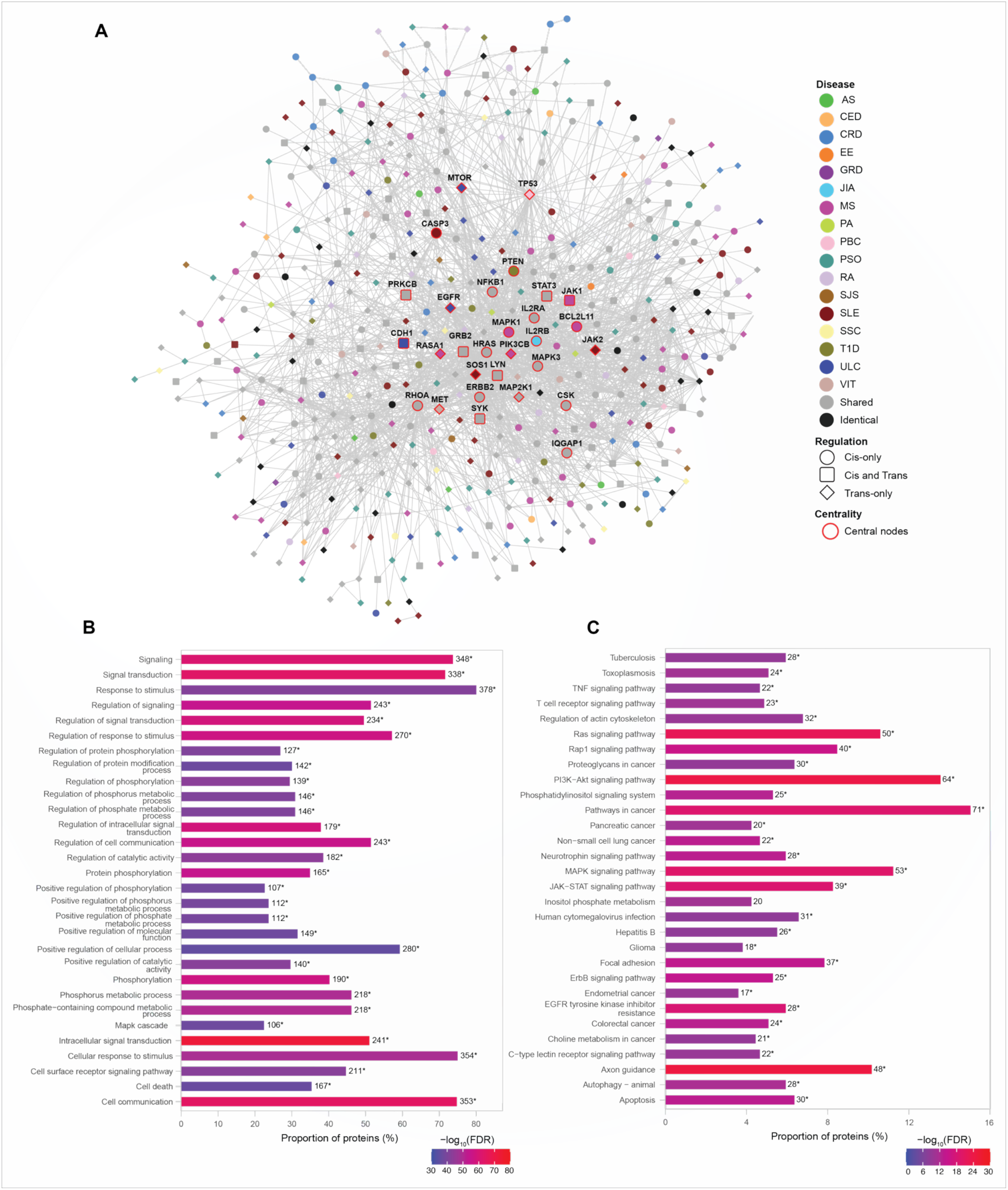
The largest network module is enriched for cellular signalling and cancer pathways. **(A)** Network representation of the module. The color of the nodes denotes the disease with which the protein is associated. Node shape indicates if the SNP acts locally (cis - circle), distally (trans - diamond), or both (cis and trans – rounded square) on the genes encoding proteins. Central nodes are highlighted in red borders and labelled. Cytoscape (version 3.8.2) was used for visualization of the module. **(B)** Module 7 is enriched for signalling and metabolic processes. The top 30 enrichment results are shown (FDR≤4.77E-36). **(C)** KEGG pathway enrichment analysis identified enrichment in signalling and cancer related pathways (FDR≤3.70E-09). The top 30 pathway enrichment results are shown. In **(B)** and **(C)**, the numbers, to the right of each bar, denote the number of proteins enriched for that term or pathway. The asterisk designates terms or pathways that were enriched for shared proteins that are central to the network (Supplementary data 12 Table 4 and 5 respectively).

## Discussion

In this study, we integrated information from different biological levels (i.e. Hi-C chromatin conformation data, eQTL data, and protein interaction data) to determine how SNPs that were independently associated with 18 AiDs might contribute to the observed multimorbidity between these conditions. Our analysis revealed a subset of genes whose transcript levels are regulated by multiple AiD-associated SNPs. We have demonstrated that these shared genes form highly connected hubs within the Ai-PPIN network, and are significantly enriched in major biological processes that include immunity, cellular metabolism and signaling cascades. The 14 highly connected modules we identified within the Ai-PPIN were significantly enriched in HLA, non-HLA, and cancer-related aspects of immunity. We contend that these observations will aid in identifying AiD specific subsets of genes that contribute to specific features of the disease and might serve as targets for drug repurposing.

The highly polymorphic HLA complex genes are among the strongest risk factors of all immune-mediated diseases. We identified 33 HLA genes that are associated with SNPs from at least two of 17 autoimmune conditions. In so doing, we provide evidence that corroborates the fundamental relevance of the HLA complex in AiDs. Notably, we did not observe any eQTL association involving HLA genes and eosinophilic esophagitis (EE) associated SNPs. This suggests that the primary risk factors for EE reside outside of the HLA genes (Kottyan et al., 2019). Despite this, the identification of eQTL SNPs for EE that regulate non-HLA genes (e.g., DOCK3, C4A, BLK, ERI1) which were also regulated by other AiDs, is evidence for the existence of a common HLA-independent genetic mechanisms for EE and other AiDs. Further support for common HLA-independent genetic mechanisms was provided by the identification of non-HLA risk loci that were associated with more than one AiD. We propose that these shared non-HLA loci contribute to variation in the immune system that alters the presentation of the driving AiD to include alternative morbidities.

Despite the incompleteness of human protein interactome maps, proteins encoded by genes associated with similar disorders show a higher likelihood of physical interactions (Goh et al., 2007). Moreover, it is widely recognized that if a gene or protein is involved in a molecular process, its direct interactors are also frequently involved in the same process (Oti, Snel, Huynen, & Brunner, 2006). Consistent with this, the proteins encoded by the genes we identified as being regulated by the AiD-associated SNPs formed highly inter-connected networks. Moreover, the functional modules we identified contained protein products encoded by genes that were subject to regulation by SNPs from between one to ten AiDs. Multiple AiD-associated SNPs regulatory impacts on these functional genetic modules is consistent with the existence of overlapping clinical presentations and common biochemical processes, or pathways. Thus, despite the apparent independence of the genetic variants that are associated with these AiDs, it is clear that the diseases are not independent at the molecular level. As such, it is likely that environmental stimulation of the pathways on which the regulatory impacts converge will initiate a cascade of events that triggers the emergence of multiple phenotypes, the severity of which is dependent upon the number of contributory genetic variants contained within individual’s genome.

The bidirectional relationship between AiDs and cancer is well-established (Giat, Ehrenfeld, & Shoenfeld, 2017). The dysregulation of genes involved in tumor suppression (e.g., *TP53*) and neoplastic processes (e.g., *ERRB2*, *EGFR*) by AiD-associated SNPs provides new insights into this complex relationship. The proteins encoded by these cancer-risk genes and other proteins encoded by AiD-associated genes were organized into a highly interconnected functional module (Module 7). Notably this module was enriched for genes associated with many cancer types (e.g., colorectal, endometrial, gastric, thyroid, breast, prostate, non-small cell lung cancer) as well as many cellular signalling (e.g., axon guidance, PI3K-Akt Ras, mTOR, MAPK signalling pathways), infectious disease (e.g., Tuberculosis, Pertussis, Influenza), and immune function (e.g., T cell receptor signalling, Th17 cell differentiation, IL-17 signalling). Collectively, these findings suggest that a subset of the AiD risk variants might increase the risk of cancer indirectly through alterations to the intermediary phenotype (i.e., gene expression) of the cancer-risk genes. It is not unreasonable to speculate that the inter-connectedness of the genes that are affected by AiD-associated SNPs, within a functional module that is enriched for cancer and immune processes, may alter the precarious balance between immune oversurveillance (AiD) and under-surveillance (unchecked growth in cancer and infectious disease) in genetically predisposed individuals.

There are a number of potential limitations to this study. Firstly, our analysis was restricted to GWAS SNPs that were identified as having both an eQTL association and physically interacting with the target genes. As such, it is possible that we have missed some proximal gene targets if they were not resolved at the level of the Hi-C restriction fragments. Secondly, most of the spatial chromatin interactions were identified from immortalized cancer cell-lines or primary tissues. By contrast, the eQTL associations were obtained mostly from post-mortem samples taken from a cross-sectional cohort (20- 70 years). Therefore, it is possible that the Hi-C interactions and eQTL sets were not representative of the tissues in which they were tested. However, in spite of this obvious technical bias, our results were reproducible and tissue-specific (FDR < 0.05) and this provide an overall systems-level understanding of the regulatory interactions observed between AiD-associated SNPs and their target genes. Thirdly, eQTL associated transcript level changes were used as a proxy for changes to gene expression. While some studies have noted a positive correlation between mRNA expression and protein expression (Schwanhüusser et al., 2011; Wilhelm et al., 2014), particularly when considering transcripts and proteins encoded by the same gene (Haiyun Wang et al., 2010), transcript-level is widely recognized as being in-sufficient to accurately predict protein levels. Despite this, these limitations should not be allowed to detract from the significance of the convergence of AiD-associated SNPs upon shared biological pathways.

In conclusion, as we move into the era of genome editing and personalized medicine, we must translate our understanding of genetic risk to the biological pathways that represent viable targets for therapeutic intervention. Our results represent one such analysis of discrete genetic data that enabled the identification of functional protein modules that putatively contribute to the shared pathogenesis underlying the development of comorbidity within AiDs. Future experiments will determine if the predictions of shared pathways will aid in the treatment of patients with multiple AiD presentations.

## URLs

GWAS catalog: https://www.ebi.ac.uk/gwas/

GTEx portal: https://www.gtexportal.org/home/

STRING database: https://string-db.org

The Drug Gene Interaction database: http://dgidb.org

## Code availability

CoDeS3D pipeline is available at https://github.com/Genome3d/codes3d-v2. Scripts used for data curation, analysis and visualization are available at https://github.com/Genome3d/Genetics_of_autoimmune_diseases Python v3.6.9 was used for all the python scripts. R v4.0.2 and RStudio v1.3.959 was used for data analyses.

## Data availability

All supplementary data is available in figshare

Supplementary data 1 - DOI: https://doi.org/10.17608/k6.auckland.14273606

Supplementary data 2 - DOI: https://doi.org/10.17608/k6.auckland.14273630

Supplementary data 3 - DOI: https://doi.org/10.17608/k6.auckland.14273633

Supplementary data 4 - DOI: https://doi.org/10.17608/k6.auckland.14273654

Supplementary data 5 - DOI: https://doi.org/10.17608/k6.auckland.14274659

Supplementary data 6 - DOI: https://doi.org/10.17608/k6.auckland.14274794

Supplementary data 7 - DOI: https://doi.org/10.17608/k6.auckland.14287652

Supplementary data 8 - DOI: https://doi.org/10.17608/k6.auckland.14288042

Supplementary data 9 - DOI: https://doi.org/10.17608/k6.auckland.14288300

Supplementary data 10 - DOI: https://doi.org/10.17608/k6.auckland.14288834

Supplementary data 11 - DOI: https://doi.org/10.17608/k6.auckland.14289158

Supplementary data 12 - DOI: https://doi.org/10.17608/k6.auckland.14287337

## Author Contributions

SG performed analyses, interpreted data, and wrote the manuscript. TF wrote CoDeS3D and commented on the manuscript. EG prepared Hi-C datasets used in the study and commented on the manuscript. WS contributed to data interpretation and commented on the manuscript. JOS directed the study, contributed to data interpretation and co-wrote the manuscript.

## Acknowledgments

This study was funded by a Ministry of Business, Innovation and Employment Catalyst grant (New Zealand- Australia LifeCourse Collaboration on Genes, Environment, Nutrition and Obesity; UOAX1611) to JO’S and SG. TF and JO’S were funded by a Health Research Council explorer grant (HRC 19/774) and a grant from the Dines Family Trust. JO’S and WS are funded by a Royal Society of New Zealand Marsden Fund (Grant 16-UOO-072). WS was also funded by a postdoctoral fellowship from the Auckland Medical Research Foundation (grant ID 1320002). EG was funded by University of Auckland Doctoral Scholarship. The authors would like to thank the Genomics and Systems Biology Group (Liggins Institute) for useful discussions. We would like to acknowledge Genotype-Tissue Expression (GTEx) consortium and the funders of GTEx Project – common Fund of the Office of the Director of the National Institutes of Health, and by NCI, NHGRI, NHLBI, NIDA, NIMH, and NINDS.

